# The magnitude of trial-by-trial neural variability is reproducible over time and across tasks in humans

**DOI:** 10.1101/096198

**Authors:** Ayelet Arazi, Gil Gonen-Yaacovi, Ilan Dinstein

## Abstract

Numerous studies have shown that neural activity in sensory cortices is remarkably variable over time and across trials even when subjects are presented with an identical repeating stimulus or task. This trial-by-trial neural variability is relatively large in the pre-stimulus period and considerably smaller (quenched) following stimulus presentation. Previous studies have suggested that the magnitude of neural variability affects behavior such that perceptual performance is better on trials and in individuals where variability quenching is larger. To what degree are neural variability magnitudes of individual subjects flexible or static? Here, we used EEG recordings from adult humans to demonstrate that neural variability magnitudes in visual cortex are remarkably consistent across different tasks and recording sessions. While magnitudes of neural variability differed dramatically across individual subjects, they were surprisingly stable across four tasks with different stimuli, temporal structures, and attentional/cognitive demands as well as across experimental sessions separated by one year. These experiments reveal that, in adults, neural variability magnitudes are mostly solidified individual characteristics that change little with task or time, and are likely to predispose individual subjects to exhibit distinct behavioral capabilities.

**Significance statement:** Brain activity varies dramatically from one moment to the next. Recent research has revealed that humans exhibit different magnitudes of trial-by-trial neural variability, which explain differences in their perceptual performance. How stable are neural variability magnitudes over time and across tasks? Here, subjects performed four different experiments in two experimental sessions separated by one year. The results revealed that neural variability magnitudes were remarkably consistent over time and across tasks, suggesting that the magnitude of neural variability is a solidified neural characteristic that may predispose individual subjects to exhibit different behavioral capabilities.

## Introduction

Neural activity in the mammalian brain is notoriously variable/noisy over time (Vreeswijk and Sompolinsky, 1996; Faisal et al., 2008). One way of studying neural variability is by quantifying trial-by-trial variability in sensory cortices across trials containing an identical stimulus. Such studies often distinguish between variability that is apparent before stimulus presentation and variability that is apparent after stimulus presentation (Arieli et al., 1996; Churchland et al., 2010; Goris et al., 2014). Recent research has shown that pre-stimulus neural variability is considerably larger than post-stimulus variability, thereby demonstrating that sensory stimulation reduces (“quenches”) ongoing neural variability (Churchland et al., 2010). Such variability quenching was reported consistently across studies examining a variety of cortical areas and arousal states, while using different types of stimuli, and when measuring neural activity with electrophysiology in animals (Monier et al., 2003; Finn et al., 2007; Mitchell et al., 2007; Churchland et al., 2010, 2011; Hussar and Pasternak, 2010) or neuroimaging in humans (He, 2013; He and Zempel, 2013; Schurger et al., 2015). An interesting exception is that during perceptual decision making, neural variability sometimes increases before the decision in made (Churchland et al., 2011).

Several lines of evidence suggest that neural variability affects behavioral performance. First, larger variability quenching in sensory cortices is associated with better perceptual performance, whether examined across trials (Schurger et al., 2015) or across individual subjects (Arazi et al., 2017). Second, actively allocating attention to a visual stimulus improves behavioral performance not only by gain modulation, but also by reducing the trial-by-trial response variability of single neurons in visual cortex (Mitchell et al., 2007) and the shared/correlated variability across the local neural population (Cohen and Maunsell, 2009; Mitchell et al., 2009). Third, increasing dopamine and norepinephrine levels increases the magnitude of neural variability in animals (Aston-Jones and Cohen, 2005; Sakata et al., 2008) and generates behavior that is more exploratory (Aston-Jones and Cohen, 2005). Fourth, it has been suggested that larger ongoing cortical variability is associated with better cognitive performance (McIntosh et al., 2008; Garrett et al., 2011, 2013).

While neural variability is under the flexible control of attention and neuromodulation to a certain extent, many of the mechanisms that generate and govern neural variability are likely to be a product of individual genetics and early development. For example, mechanisms that govern the reproducibility of neural activity by maintaining stable excitation-inhibition balances (Turrigiano, 2011) and reliable synaptic transmission (Ribrault et al., 2011), are the product of individual genetics and environmental exposure during early critical periods (Hensch, 2005; Takesian and Hensch, 2013). Since individual subjects have different genetics and experience different environments, one may expect intrinsic neural variability magnitudes to differ across individuals and potentially predispose them to different behavioral capabilities.

With this in mind, we hypothesized that neural variability magnitudes of individual adult subjects are, to a large extent, a solidified neural characteristic. A prediction of this hypothesis is that individual subjects should exhibit distinct magnitudes of neural variability that would be reproducible across experiments with different stimuli, tasks, and over time. To test this, we measured neural variability in visual cortex with EEG while subjects performed four tasks that differed in their structure, stimulus, attentional demands, and cognitive requirements. The same subjects performed all four experiments in two experimental sessions separated by a year. We then quantified the neural variability magnitudes of individual subjects in each of the experiments and experimental sessions to determine their consistency across experiments and over time.

## Materials and Methods

### Subjects

Twenty-four subjects (eight males, mean age during the first session= 23.7 years, SD= 1.4) took part in this study. All subjects had normal or corrected-to-normal vision. The study was approved by the Ben-Gurion University Internal Review Board. Subjects provided written informed consent during both experimental sessions and were either paid for their participation or received research credit.

### Experimental design

All subjects completed four experiments in each of the two experimental sessions. The gap in time between the first and the second session was 12.3 months on average (SD = 1.1). The study was performed in a dark and sound proof room. The stimuli were presented using MATLAB (Mathworks, Inc., USA) and Psychtoolbox (Brainard, 1997).

#### Checkerboard experiment

The visual stimulus consisted of a checkerboard annulus with an inner radius of 0.6° visual angle and an outer radius of 3.7° visual angle. The experiment contained 600 trials: 400 trials with the stimulus and 200 trials where the stimulus was omitted. The stimulus was presented for 50ms and followed by a randomized inter-trial interval lasting 750-1200ms. The experiment also included an orthogonal color-detection task at fixation, which was intended to divert attention away from the checkerboard stimulus. Subjects were instructed to press a key whenever the black fixation cross changed its color to gray. The experiment contained 80 random color changes, which lasted 30ms and subjects had one second to respond. Correct and incorrect responses were indicated by changing the fixation cross to green or red, respectively.

#### Choice reaction time (CRT) experiment

A black triangle or a circle was presented at the center of the screen for 300ms on each trial. Subjects were instructed to press the right or left arrow keys, respectively, as quickly as possible using their right index finger. Each trial was followed by an inter-trial interval of 1200ms. A total of 200 trials were presented, 100 trials with each of the two stimuli.

#### Go-no-go experiment

Stimuli and structure were identical to those described in the CRT experiment, except that participants were instructed to press the spacebar as quickly as possible with their right index finger whenever they saw a circle (“go” trial) and not when the triangle was presented (“no go” trial). A total of 300 trials were presented and 80% of the trials contained the “go” stimulus.

#### 2-back experiment

Stimuli were composed of 4 Chinese letters, presented at the center of the screen and participants were asked to press the "J" key whenever the current letter matched the one that was presented 2 trials before. Each letter was presented for 500ms and followed by an inter trial interval of 500ms. A total of 300 trials were presented with third of them containing a 2-back repeat.

### EEG and eye tracking recordings

EEG data were recorded using a 64-channel BioSemi system (Biosemi Inc., Netherlands), connected to a standard EEG cap according to the international 10-20 system. Data were referenced to the vertex electrodes. Electrooculography (EOG) was recorded using two electrodes at the outer canthi of the left and right eyes and one electrode placed below the right eye. In the Checkerboard experiment, the position of the right eye was recorded using an eye-tracker (EyeLink 1000, SR-research) at a sampling rate of 1000Hz.

### EEG preprocessing

Data was analyzed using Matlab and the EEGLAB toolbox (Delorme and Makeig, 2004). Continuous EEG data was down sampled to 512Hz, filtered using a 1-40 Hz band pass filter, and re-referenced to the bilateral mastoid electrodes. EEG epochs were extracted using a time window of 700ms (200ms pre-stimulus to 500ms post-stimulus) and baseline correction was not performed so as not to alter trial-by-trial variability in the pre-stimulus interval. In the Checkerboard experiment only trials where stimulus was presented were extracted, in the CRT experiment trials with both stimuli (circle or triangle) were extracted, in the go-no-go experiment only the “go” trials were extracted and in the 2-back experiment trials with the four different stimuli (Chinese letters) were extracted.

Epochs containing absolute amplitudes that exceeded 70μV or where the power exceeded 25db in the 20-40Hz frequency range were identified as containing eye blinks or muscle artifacts, respectively, and were removed from further analysis. In the Checkerboard experiment identification of eye blinks was confirmed by eye tracking; trials containing horizontal or vertical eye movements that exceeded 1.5 SD of the mean were identified as trials where fixation was not maintained (i.e. trials containing saccades) and excluded from EEG analyses. Mean number of trials across subjects and sessions after trials rejection in the four experiment was 249 trials in the Checkerboard experiment (SD=50), 146 trials in the CRT experiment (SD=39), 161 trials in the go-no-go experiment (SD=53), and 254 trials in the 2-back experiment (SD=46). The mean number of trials did not differ between the first and second experimental sessions.

### EEG data analysis

Trial by trial variability was computed for each time-point in the extracted epochs (-200 to 500ms) for each of the 64 electrodes, in each subject separately. Trials from the first and second experimental sessions were analyzed separately. Absolute level of trial-by-trial variability in the pre-stimulus interval was computed within a time window of −200ms and 0ms pre-stimulus – the variance was computed across trials for each time-point and then averaged across the time-points within the window. Absolute level of trial-by-trial variability in the post-stimulus interval was computed in the same manner within a time window of 150-400ms post-stimulus.

Relative trial-by-trial variability was computed by converting the variability time courses to percent change units relative to the mean trial-by-trial variability in the pre-stimulus period (-200 to 0ms). We then estimated variability quenching for each subject in each task and session by computing the difference in variability between the pre-stimulus period (-200 to 0ms) and post-stimulus period (150 to 400ms). We focused our analyses on the four occipital electrodes (O1, O2, PO7 and PO8) with the strongest visual responses.

To ensure that changes in variability were not driven by changes in the mean EEG activity, we also performed a control analysis, where we computed the coefficient of variation (CV) by dividing the magnitude of variability by the area under the curve of the mean ERP response (i.e., the ERP amplitude). This was computed separately for the pre (-200 to 0ms) or post (150 to 400ms) stimulus intervals. We then computed CV quenching as the relative change in the CV between the pre and post-stimulus periods (in units of percent change). To examine the temporal dynamics of the CV, we used a sliding window with a width of 50ms and overlap of 5ms (Figure 6).

### Behavioral data analysis

Mean accuracy, mean reaction time (RT) and reaction time variability (across trials) was computed for each subject and each session, in CRT, go-no-go and two-back tasks as well as the color-detection task in the Checkerboard experiment. The first 10 trials in each experiment and trials with RT below 200ms were excluded from the analysis. Trials with incorrect responses were excluded from the RT analyses.

### Statistical Analysis

We assessed relationships across measures using Pearson’s correlations. The statistical significance of the correlation coefficients was assessed with a randomization test where we shuffled the labels of the subjects before computing the correlation coefficient. This procedure was performed 10,000 times while shuffling the labels across subjects randomly each time to generate a null distribution for each pair of EEG/behavioral measures. For the true correlation coefficient to be considered significant it had to exceed the 97.5^th^ percentile or lower than the 2.5^th^ percentile of this null distribution (i.e., equivalent to a p-value of 0.05 in a two-tailed t-test). Comparisons across experiments/tasks were performed using a one-way ANOVA with task as the only factor, followed by post hoc Tukey’s tests when the initial result indicated significant differences. When performing correlations between neural variability and behavioral measures for each of the 64 electrodes, we used the false discovery rate (FDR) correction (Benjamini and Hochberg, 1995) to control for the multiple comparisons problem (Figure 5).

### Electrode offset variability

The Biosemi EEG system utilizes active electrodes, which do not yield a measure of impedance. Instead, fluctuations in electrode offset (i.e. slow changes in the voltage potential over time) are considered the best indication for the quality of EEG recording (Kappenman and Luck, 2010). We, therefore, computed the electrode offset variability across trials for each subject during each of the experiments in each experimental session. We computed the offset value for each trial, the variability across trials in each of the four examined electrodes, and finally the mean across electrodes in each experiment. We then correlated offset variability with the EEG variability measures to check if differences in the quality of EEG recordings across individuals could explain our results.

### Gaze variability

Gaze position was measured during the Checkerboard experiment only. We computed the distance from the fixation cross at each time point from stimulus onset to 500ms post-stimulus, then computed the standard deviation across trials for each time point, and finally averaged across all time points (0-500ms) to generate a single measure of gaze variability per subject. We correlated gaze variability across the first and second sessions to determine whether individual subjects exhibited reproducible gaze variability across sessions. Three subjects were excluded from this analysis due to difficulties in the calibration process of the eye tracker in one of sessions.

## Results

Twenty-four subjects completed two experimental sessions separated by one year. Each session included four experiments that differed in their structure, stimulus, attentional demands, and cognitive loads. In the first experiment, subjects passively observed a checkerboard annulus on each trial, while their task was to identify and report infrequent color changes of the fixation cross. This enabled us to measure neural variability magnitudes to an un-attended stimulus (i.e., the checkerboard). In the second experiment, subjects performed a choice reaction time (CRT) task where they responded with one button to a circle stimulus and with another button to a triangle stimulus. This enabled us to measure neural variability magnitudes to an attended stimulus during a very easy task. In the third experiment, subjects performed a go-no-go task where they responded only to the circles (go trials) and not to the triangles (no-go trials). This enabled us to measure neural variability magnitudes to an attended stimulus during a somewhat harder task that required response inhibition. In the final experiment, subjects performed a 2-back task where they were presented with alternating Chinese letters and instructed to respond whenever the current letter matched the letter that was presented two trials before. This enabled us to measure neural variability magnitudes to an attended stimulus during a difficult working memory task.

Differences in the attentional and cognitive demands of the four tasks were clearly evident in the behavioral performance of the subjects (Figure 1). One-way ANOVA analyses demonstrated that there were clear differences in the accuracy rates (*F*_(3,92)_ = 89.2, *p* = 0.4 × 10^−26^), mean reaction times (*F*_(3, 92)_ = 56.3, *p* = 0.9x10^−20^), and reaction time variability (F_(3, 92)_ = 106.8, p = 0.7x10^−29^) across the four tasks. Post-hoc Tukey’s tests revealed that there were significant differences across all pairs of tasks (*p* < 0.01 for all behavioral measures), except for the CRT and Go-no-go tasks. Specifically, accuracy rates were higher, mean reaction times were lower, and reaction time variability was lower in the CRT and Go-no-go tasks as compared with the color-detection task in the Checkerboard experiment and the 2- back task. In addition, accuracy rates were significantly higher, mean reaction times were lower, and reaction time variability was lower in the color-detection task as compared with the 2-back task. This demonstrates that the CRT and Go-no-go tasks were easier than the color-detection and 2-back tasks and that the 2-back task was harder than the color-detection task.

**Figure 1:**
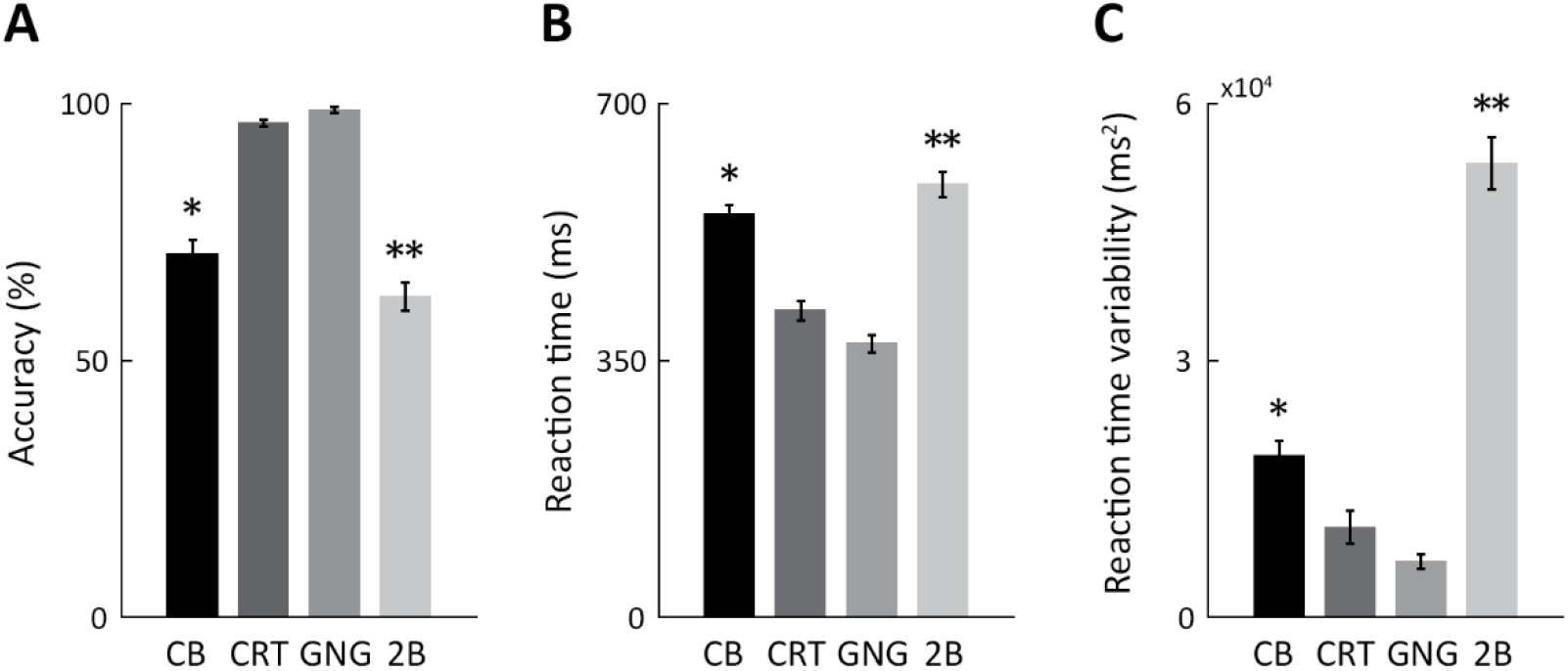
Behavioral performance measures. Mean across subjects and sessions for accuracy (A), reaction time (B) and reaction time variability (C) in each of the four tasks. Error bars: standard error of the mean across subjects. Asterisks: Significant differences across experiments (Post-hoc Tukey’s tests, *p* < 0.01). One asterisk: Significant differences between CB experiment and CRT or GNG experiments. Two asterisks: Significant differences between 2B experiment and all other experiments. CB: checkerboard, CRT: choice reaction time, GNG: go-no-go, 2B: 2-back.

Note that in the Checkerboard experiment, the relatively difficult color-detection task diverted the subjects’ attention away from the checkerboard stimulus, thereby allowing us to quantify trial-by-trial neural variability to an unattended stimulus. In contrast, the 2-back task required that subjects attend the stimulus, thereby allowing us to quantify trial-by-trial neural variability to a strongly attended stimulus.

### Neural variability quenching

We examined trial-by-trial neural variability as a function of time before and after stimulus presentation in each of the four experiments (Figure 2). Trial-by-trial variability was reduced (i.e., quenched) following stimulus presentation in all experiments and in both recording sessions performed a year apart. Variability quenching was sustained from 150 to 400ms after stimulus presentation and most evident in occipital electrodes (O1, O2, PO7 and PO8). We quantified variability quenching as the relative change (in units of percent change) between pre-stimulus (-200 to 0ms) and post-stimulus (150 to 400ms) periods, while focusing our analyses on the four electrodes noted above. Note that we filtered our EEG data using a bandpass filter (1-40 Hz, see Materials and Methods) as commonly performed in many EEG experiments. This means that the quantified trial-by-trial variability was generated by EEG activity in this frequency range and mostly dominated by lower frequencies.

**Figure 2:**
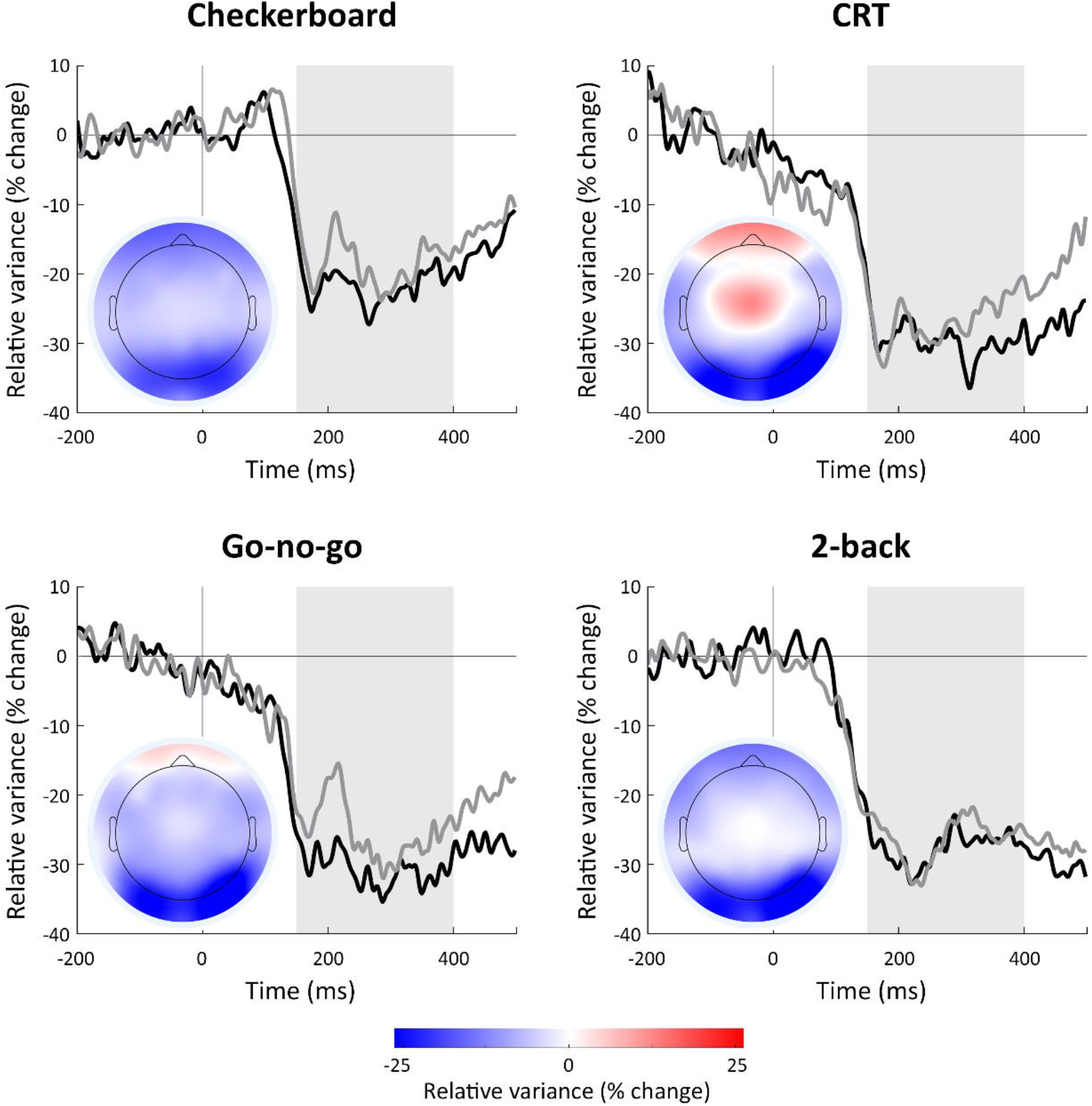
Temporal and spatial dynamics of trial-by-trial neural variability. Each time-course represents the changes in relative trial-by-trial variability (percent-change units relative to the pre-stimulus period, mean across the 4 selected electrodes) during the first (black) or second (grey) experimental session, which were separated by one year. Each panel displays the results of a different experiment. Gray background: 150-400ms post-stimulus period with sustained variability quenching that was selected for further analyses. Insets: topographic maps of variability quenching magnitudes during the 150-400ms window, demonstrating that quenching was strongest in occipital electrodes across all four experiments.

### Neural variability is a stable neural characteristic

We quantified three measures of trial-by-trial variability for each subject. Absolute trial-by-trial variability was quantified in the pre-stimulus (-200 to 0ms) and poststimulus (150-400ms) periods for each subject, in each of the four experiments, and each of the experimental sessions (see Materials and Methods). Variability quenching was quantified as the difference between variability magnitudes in the pre and post stimulus periods. All three measures of variability were strongly and significantly correlated across the two EEG recording sessions in each of the four experiments (*r*(24) > 0.58, *p* < 0.003, Figure 3). This demonstrates that the neural variability magnitudes of individual subjects barely changed over a one year period.

**Figure 3:**
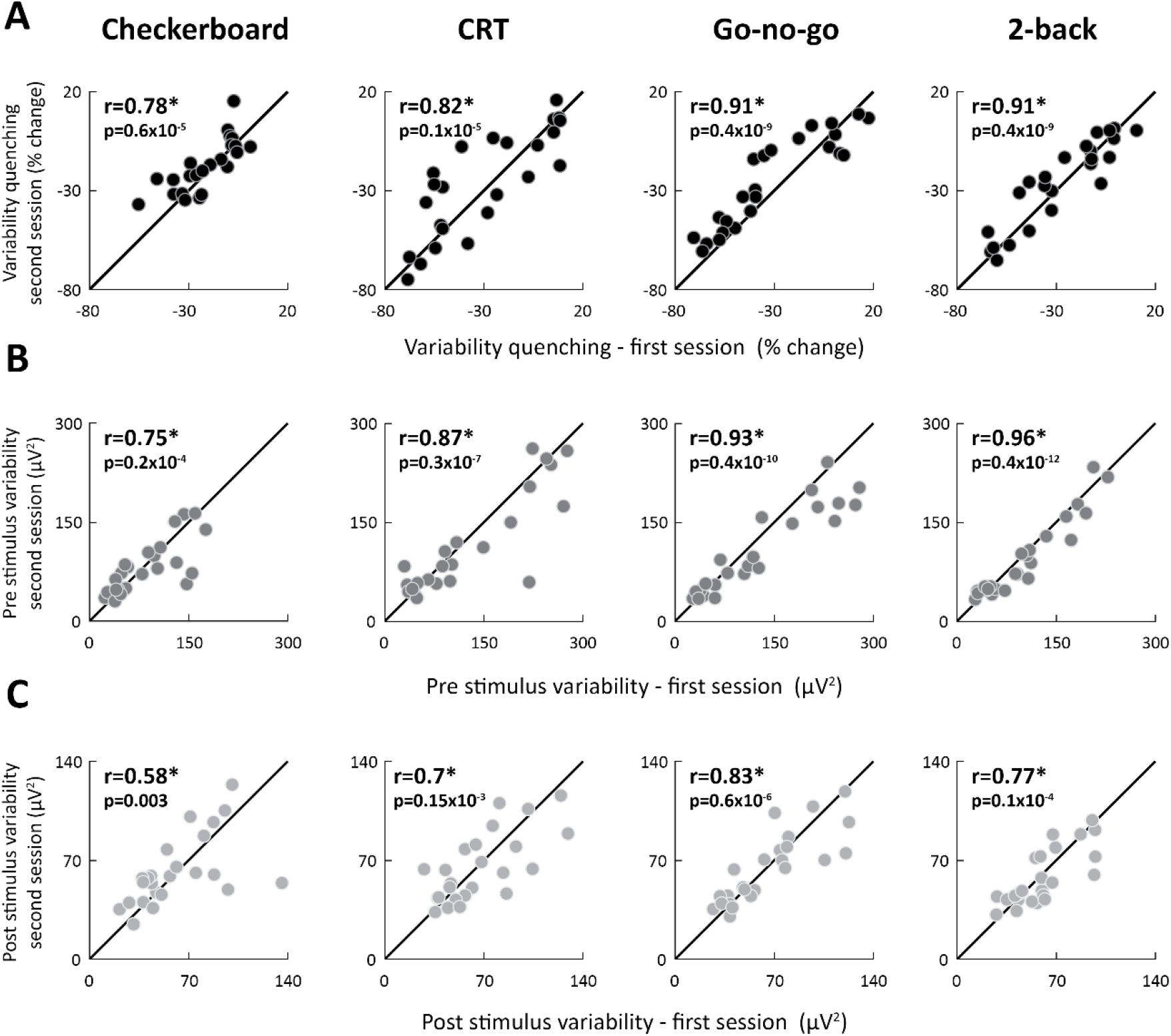
Individual neural variability magnitudes were consistent across experimental sessions separated by one year. Scatter plots present the magnitudes of variability quenching (A), pre-stimulus variability (B), and post-stimulus variability (C) in individual subjects during the first and second experimental sessions for each of the four experiments. The unity line is drawn for reference in each panel. Each point represents a single subject. Asterisks: significant correlation as assessed by a randomization test (*p* < 0.003). Pearson’s correlation coefficients and p-values are noted in each panel.

Individual variability magnitudes were also strongly correlated across experiments despite their distinct stimuli, temporal structures, and attentional/cognitive demands. Given the strong correlations across sessions (Figure 3), we averaged each of the variability measures across the two sessions. We then compared individual variability magnitudes across experiment pairs. This analysis revealed strong, positive, and significant correlations across all pairs of experiments when examining variability quenching (*r*(24) > 0.74, *p* < 0.4 *x* 10^−4^, Figure 4), pre-stimulus variability (*r*(24) > 0.86, *p* < 0.8 *x* 10 ^7^, Table 1), or post-stimulus variability (*r*(24) > 0.89, *p* = 0.4 *x* 10^−8^, Table 1) magnitudes.

**Figure 4:**
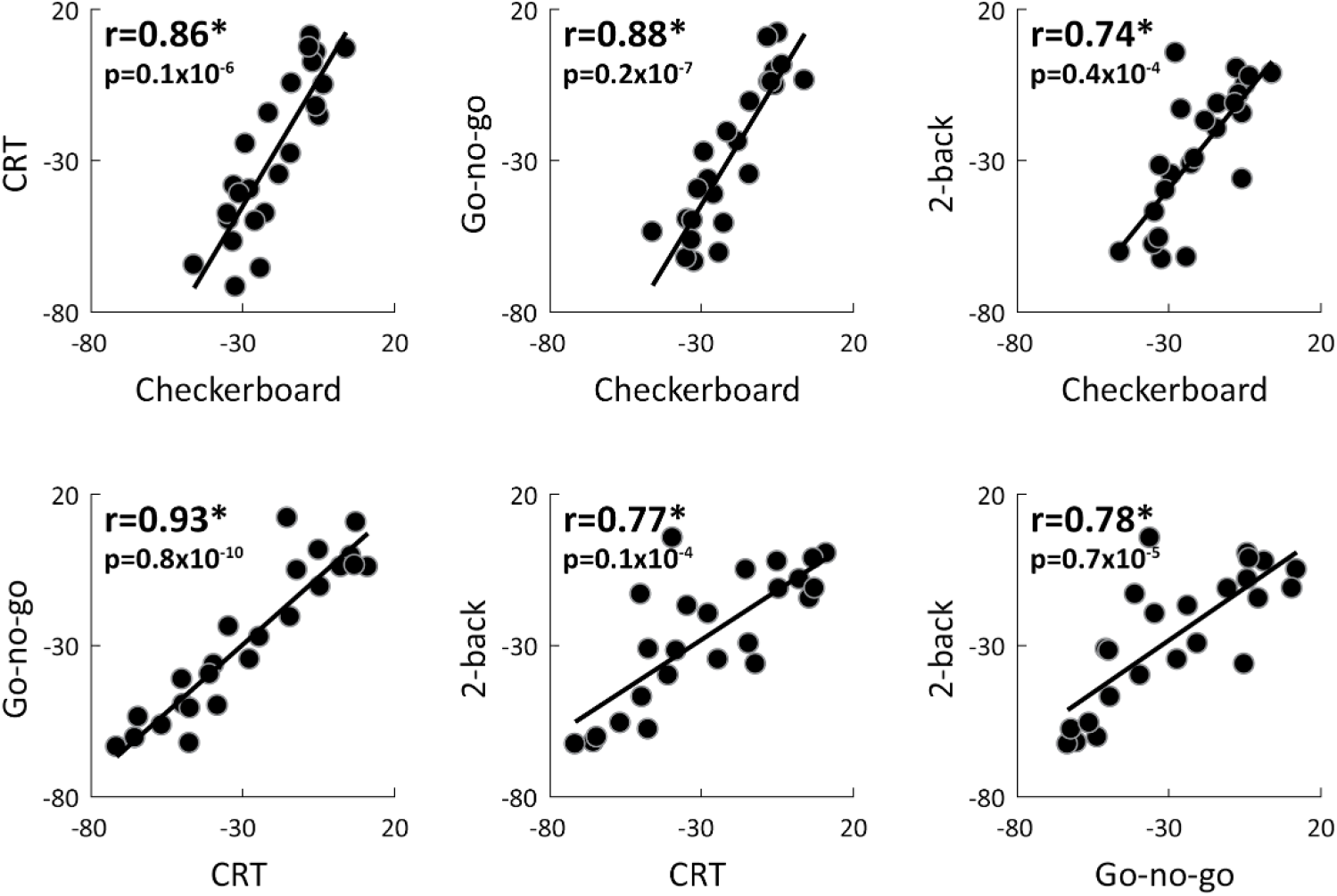
Individual variability quenching magnitudes were consistent across experiments. Scatter plots demonstrate the relationship between variability quenching magnitudes in each pair of experiments. Each dot represents a single subject. The linear fit is drawn for reference in each panel. Asterisks: significant correlation as assessed by a randomization test (*p* < 0.4 *x* 10^−4^). Pearson’s correlation coefficients are noted in each panel.

**Table 1:** individual magnitudes of pre-stimulus (top row) and post-stimulus (bottom row) neural variability were strongly correlated across experiments. Pearson’s correlation coefficients and p-values are noted for each pair of experiments. CB: checkerboard, CRT: choice reaction time, GNG: go-no-go, 2B: 2-back.

|  | CB-CRT | CB-GNG | CB-2B | CRT-GNG | CRT-2B | GNG-2B |
| --- | --- | --- | --- | --- | --- | --- |
| Pre stimulus | $r = 0.86$<br>$p = 0.5 \times 10^{-7}$ | $r = 0.93$<br>$p = 0.8 \times 10^{-10}$ | $r = 0.86$<br>$p = 0.8 \times 10^{-7}$ | $r = 0.96$<br>$p = 0.3 \times 10^{-13}$ | $r = 0.89$<br>$p = 0.6 \times 10^{-8}$ | $r = 0.93$<br>$p = 0.8 \times 10^{-10}$ |
| Post stimulus | $r = 0.9$<br>$p = 0.1 \times 10^{-8}$ | $r = 0.9$<br>$p = 0.2 \times 10^{-8}$ | $r = 0.89$<br>$p = 0.4 \times 10^{-8}$ | $r = 0.95$<br>$p = 0.1 \times 10^{-11}$ | $r = 0.9$<br>$p = 0.1 \times 10^{-8}$ | $r = 0.94$<br>$p = 0.1 \times 10^{-10}$ |

### Neural variability and behavioral performance

We examined the relationship between individual measures of neural variability and behavioral performance by computing the correlations between each of the three behavioral measures (accuracy rate, reaction time and reaction time variability) and each of the three variability measures (pre-stimulus variability, post-stimulus variability and variability quenching). No significant correlations were found when we separately examined data from the first (Accuracy: −0.13 < *r*(24) < 0.22, *p* > 0.31; RT: −0.28 < *r*(24) < 0.23, *p* > 0.18; RT variability: −0.34 < *r*(24) < 0.3, *p* > 0.1) or second (Accuracy: −0.32 < *r*(24) < 0.28, *p* > 0.12; RT: −0.13 < *r*(24) < 0.31, *p* > 0.13; RT variability: −0.27 < *r*(24) < 0.26, *p* > 0.2) experimental sessions, nor when we computed the mean behavioral\variability measures across sessions (Table 2).

**Table 2:** Relationship between measures of neural variability and behavioral measures. Pearson’s correlation coefficients and p-values for each behavioral (accuracy, mean RT or RT variability) and each variability (Quenching, pre-stimulus or post-stimulus) measure. Task abbreviations, CB: checkerboard, CRT: choice reaction time, GNG: go-no-go, 2B: 2-back.

|  | Accuracy |  |  | Mean RT |  |  | RT variability |  |  |
| --- | --- | --- | --- | --- | --- | --- | --- | --- | --- |
|  | Quench | Pre | Post | Quench | Pre | post | Quench | Pre | Post |
| CB | $r=0.17$<br>$p=0.43$ | $r=-0.21$<br>$p=0.3$ | $r=-0.12$<br>$p=0.57$ | $r=0.12$<br>$p=0.55$ | $r=-0.05$<br>$p=0.8$ | $r=0$<br>$p=0.98$ | $r=-0.02$<br>$p=0.92$ | $r=-0.06$<br>$p=0.77$ | $r=-0.13$<br>$p=0.54$ |
| CRT | $r=0.13$<br>$p=0.55$ | $r=-0.05$<br>$p=0.82$ | $r=0.07$<br>$p=0.75$ | $r=0.14$<br>$p=0.5$ | $r=-0.05$<br>$p=0.8$ | $r=0.17$<br>$p=0.4$ | $r=-0.1$<br>$p=0.63$ | $r=0.2$<br>$p=0.35$ | $r=0.22$<br>$p=0.29$ |
| GNG | $r=-0.03$<br>$p=0.88$ | $r=-0.15$<br>$p=0.48$ | $r=-0.18$<br>$p=0.38$ | $r=0.08$<br>$p=0.7$ | $r=0.04$<br>$p=0.86$ | $r=0.17$<br>$p=0.41$ | $r=0.02$<br>$p=0.93$ | $r=0.14$<br>$p=0.5$ | $r=0.29$<br>$p=0.16$ |
| 2B | $r=-0.13$<br>$p=0.54$ | $r=0.13$<br>$p=0.53$ | $r=0.21$<br>$p=0.32$ | $r=-0.2$<br>$p=0.35$ | $r=0.1$<br>$p=0.62$ | $r=0$<br>$p=0.96$ | $r=-0.36$<br>$p=0.09$ | $r=0.22$<br>$p=0.3$ | $r=0$<br>$p=0.99$ |

In an additional analysis we examined these relationships separately for each electrode and present the results as a spatial correlation map (Figure 5). The spatial correlation maps differed across experiments, suggesting a lack of robust relationship between neural variability and behavioral measures in any of the electrodes. Furthermore, there were no significant correlations in any of the electrodes following FDR correction.

**Figure 5:**
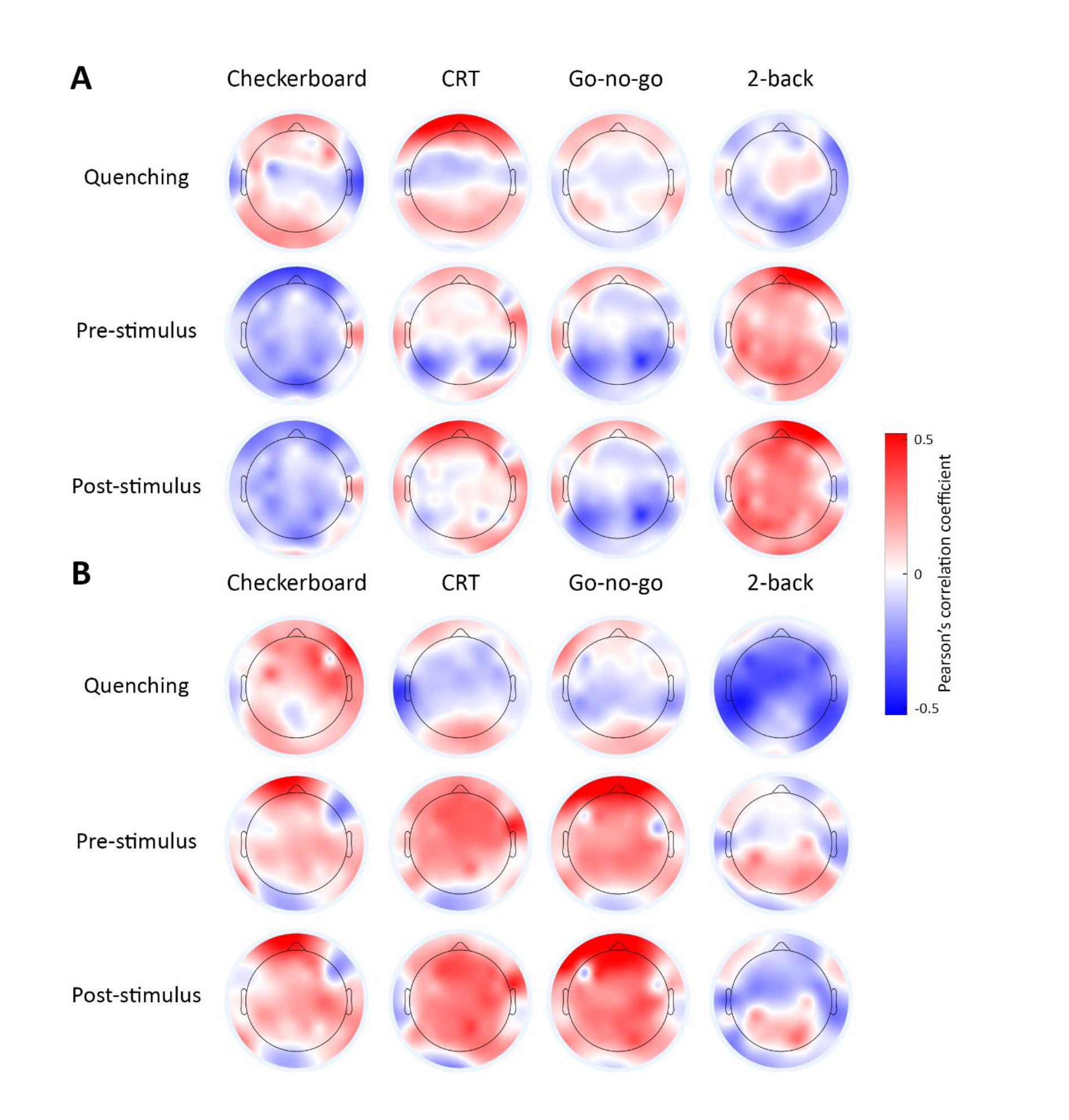
Scalp maps representing the correlation between measures of neural variability (quenching, pre-stimulus, or post-stimulus) and behavioral measures: accuracy (A) or RT (B). Color bar: Magnitude of Pearson’s correlation coefficients.

### Signal strength and trial-by-trial variability

Previous studies have demonstrated that trial-by-trial variability is associated with the mean strength of the neural response (Churchland et al., 2010). To demonstrate that the findings described above are independent of between-subject differences in the mean EEG response amplitudes, we also performed the analysis using the coefficient of variation (CV: trial-by-trial variability divided by the mean EEG response, see Materials and Methods). As with trial-by-trial variability (Figure 2), the CV also exhibited a strong reduction following stimulus presentation, within the same time-window, across all four experiments (Figure 6).

**Figure 6:**
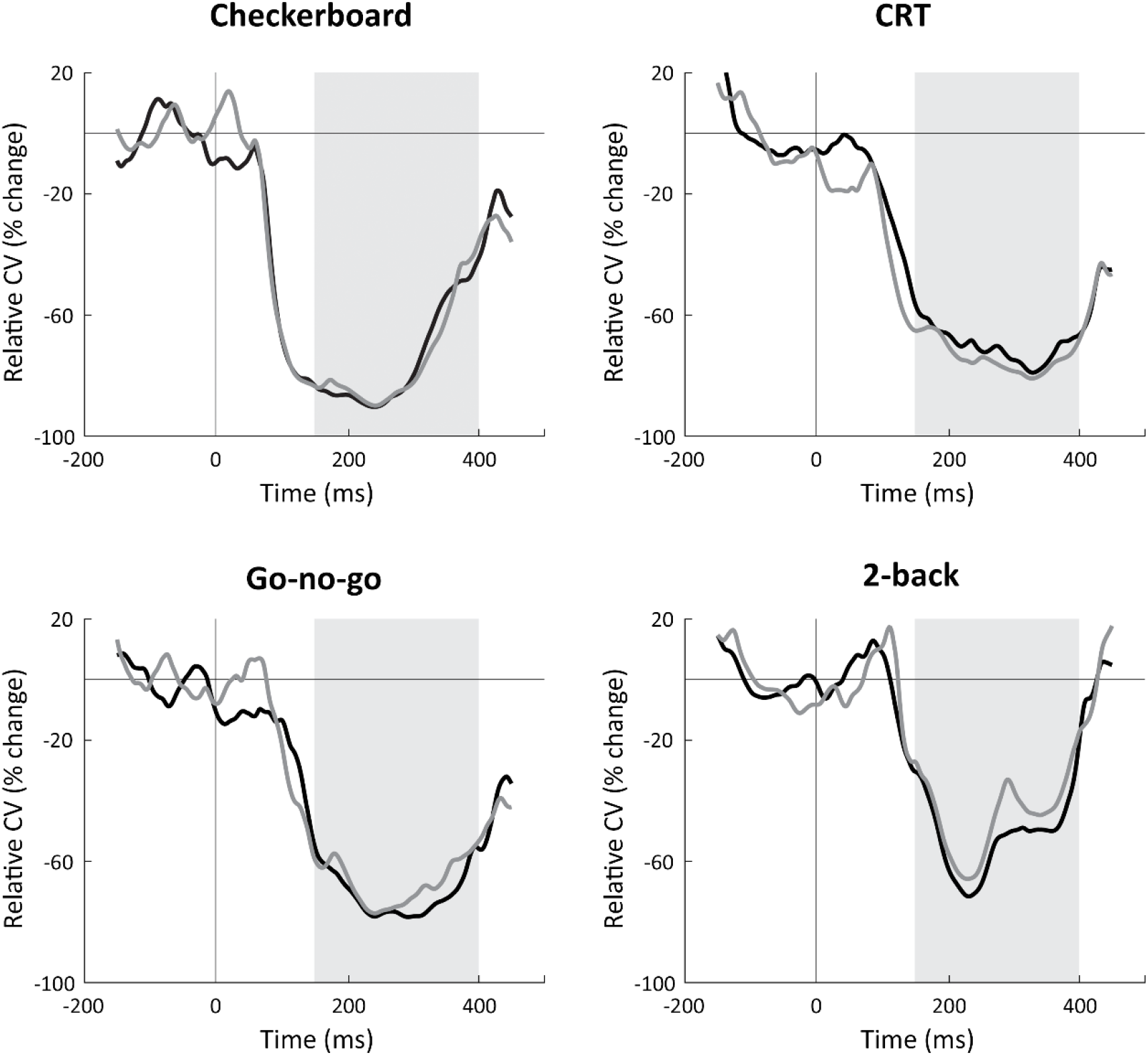
Temporal dynamics of the coefficient of variation (CV) in percent change units relative to pre-stimulus period. Each panel presents results from a single experiment in the first (black) and second (gray) experimental sessions. Gray background: time window (150-400ms) of sustained variability quenching that was selected for the previous analyses.

Most importantly, the CV magnitudes of individual subjects exhibited significant positive correlations across sessions (Quenching: 0.61 < *r*(24) < 0.82, pre-stimulus: 0.48 < *r*(24) < 0.77, post-stimulus: 0.59 < *r*(24) < 0.76; *p* < 0.018) and across most of the tasks (Quenching: 0.44 < *r*(24) < 0.91, pre-stimulus: 0.41 < *r*(24) < 0.66, post stimulus: 0.43 < *r*(24) < 0.78; *p* < 0.048).

Exceptions included the following: Checkerboard and CRT (pre-stimulus: *r*(24) = 0.26, *p* = 0.22; post-stimulus: *r*(24) = 0.26, *p* =0.22), Checkerboard and Go-no-go (post-stimulus: *r*(24) = 0.15, *p* = 0.49), and Checkerboard and 2-back (pre-stimulus: *r*(24) = 0.36, *p* = 0.08). Note that even in these cases, correlations were always positive and most were close to significant. Individual magnitudes of CV were, therefore, entirely consistent over time and mostly consistent across tasks.

### Alternative sources of trial-by-trial variability

We examined whether non-neural sources of variability, such as gaze variability or the quality of EEG recordings could explain our results regarding consistency across tasks or sessions. We utilized eye tracking data from the checkerboard experiment to demonstrate that individual magnitudes of neural variability quenching were not significantly correlated with gaze position variability in either the first (*r*(21) = −0.16, *p* = 0.49, uncorrected) or second (*r*(21) = −0.29, *p* = 0.2, uncorrected) recording session. Furthermore, re-running our analysis after regressing out individual measures of gaze position variability revealed equivalent results to those described above (Figure 3&4). Specifically, variability magnitudes remained correlated over time (Quenching: *r*(21) = 0.78, *p* = 0.3 *x* 10^−4^; pre-stimulus: *r*(21) = 0.76, *p* = 0.6 *x* 10^−4^; post-stimulus: *r*(21) = 0.6, *p* = 0.0036). This reassured us that the consistent magnitudes of neural variability described above were not associated with the ability of the subjects to maintain fixation.

To demonstrate that our results were not due to individual differences in the quality of the EEG recordings, we computed the electrode offset variability (see Materials and Methods). Electrode offset variability, was not significantly correlated with the magnitude of variability quenching in any of the experiments performed in either the first or second session (−0.25 < *r*(24) < 0.24, *p* > 0.24, uncorrected).

Furthermore, re-running our analysis after regressing out individual magnitudes of electrode offset variability revealed equivalent results to those described above (Figure 3&4). Specifically, variability magnitudes remained correlated over time (quenching: 0.8 < *r*(24) < 0.89, pre-stimulus: 0.76 < *r*(24) < 0.92, post-stimulus: 0.59 < *r*(24) < 0.74; *p* < 0.0025) and across tasks (quenching: 0.57 < *r*(24) < 0.93, pre-stimulus: 0.73 < *r*(24) < 0.96, post-stimulus: 0.76 < *r*(24) < 0.94, *p* < 0.004).

## Discussion

Our results demonstrate that neural variability magnitudes differ across adult subjects in a consistent and reproducible manner over long periods of time and across tasks with different stimuli, structures, and attentional/cognitive demands. This was true for neural variability magnitudes in either pre-stimulus or poststimulus periods and for variability quenching magnitudes (Figures 3&4, Table 1). These results suggest that neural variability magnitudes in sensory cortices are mostly stable individual characteristics that are modulated to a relatively small degree by task demands. While these reliable individual differences in neural variability magnitudes were not associated with cognitive performance measures in the current study (Figure 5), they have previously been associated with perceptual performance measures in several studies using different tasks and neuroimaging techniques (Schurger et al., 2010, 2015; Xue et al., 2010; Arazi et al., 2017). These stable neural variability magnitudes may, therefore, predispose individual subjects to exhibit distinct behavioral capabilities.

### Flexibility of neural variability

To what degree is neural variability under flexible behavioral control? Previous electrophysiology studies have reported that visual attention reduces trial-by-trial response variability while improving the accuracy of behavioral performance (Mitchell et al., 2007, 2009; Cohen and Maunsell, 2009). In contrast, raising the levels of dopamine and/or norepinephrine increases the magnitude of ongoing neural variability (Aston-Jones and Cohen, 2005; Sakata et al., 2008; Garrett et al., 2015). This may generate behavioral states associated with exploration, where the subject behaves in a more variable manner that enables learning through trial-and-error (Aston-Jones and Cohen, 2005).

While attention and neuromodulation are invaluable mechanisms for flexibly changing the magnitude of neural variability and consequent behaviors, individual differences in neural variability magnitudes are also governed by many other neurophysiological mechanisms including the noisy response characteristics of peripheral sensors (Schneeweis and Schnapf, 1999), the stochastic nature of synaptic transmission (Ribrault et al., 2011), and the dynamic changes caused by neural adaptation (Clifford et al., 2007) and synaptic plasticity (Feldman, 2009). Additional neural variability is generated by adjustments of the excitation/inhibition balance (Turrigiano, 2011) and continuous interaction and competition across large neural populations (Kelly et al., 2008). These mechanisms are likely to be the product of genetic and environmental factors that govern the development and solidification of neural circuits during critical periods (Hensch, 2005).

We believe that our results highlight the role of these later mechanisms in creating and maintaining stable individual differences in neural variability. While attention and neuromodulation are critical for adaptive behaviour, we speculate that their ability to alter the neural variability magnitude of an adult individual is relatively small and limited by these static intrinsic mechanisms. Future research could further quantify the flexibility of neural variability in individual subjects and determine whether some individuals are more flexible than others. Furthermore, studying measures of neural variability in young children would be particularly interesting for understanding how neural variability changes during development. Analogous behavioral research in humans (MacDonald et al., 2006) and birds (Ölveczky et al., 2011) has already shown that behavioral variability diminishes during early development and stabilizes in adulthood.

### The behavioral significance of neural variability

There is ongoing debate regarding the potential behavioral significance of different measures of neural variability. On the one hand, several studies have demonstrated that smaller trial-by-trial neural variability is associated with better perceptual performance and memory encoding. For example, fMRI studies have reported that trial-by-trial variability is smaller on trials where a threshold-level stimulus is detected (Schurger et al., 2010) and on trials where a stimulus is later remembered (Xue et al., 2010). Similarly, MEG and EEG studies have reported that neural variability quenching in sensory cortices is larger on trials where a threshold-level stimulus is detected (Schurger et al., 2015) and in individuals with lower (better) contrast discrimination thresholds (Arazi et al., 2017). Faster reaction times were apparent in trials with smaller trial-by-trial variability in firing rates of macaque V4 neurons (Steinmetz and Moore, 2010), smaller trial-by-trial fMRI variability (He, 2013) and in trials where ECOG activity that was more similar to the mean response across trials (He and Zempel, 2013). Furthermore, excessive neural variability in sensory cortices has been reported in different disorders including autism (Milne, 2011; Dinstein et al., 2012), ADHD (Gonen-Yaacovi et al., 2016), and schizophrenia (Yang et al., 2014), while electrophysiology studies have reported that neural responses are more variable in elderly animals (Turner et al., 2005; Yang et al., 2009) and humans (Anderson et al., 2012) who exhibit cognitive decline. These results are in line with signal detection theory principles, which suggest that intrinsic variability/noise reduces the detection and discrimination abilities of a perceptual system (Green and Swets, 1966).

Other studies, however, have reported that younger individuals exhibit larger fMRI time-course variability than elderly individuals (Garrett et al., 2010) and that this coincides with faster and more consistent responses when performing cognitive tasks such as perceptual matching, attentional cueing, and delayed match to sample (Garrett et al., 2013). It has been proposed that such increased ongoing variability may be beneficial for cognitive performance, because it allows for higher neural complexity and enables a neural network to flexibly switch between different states (McIntosh et al., 2008). Note that certain tasks, such as perceptual decision making, may generate an increase in neural variability before the decision is made (Churchland et al., 2011). In our study, we note that neural variability increased following stimulus presentation in the CRT task in fronto-central electrodes (Figure 2). The specificity of this finding to the CRT task and its potential cognitive significance remain to be determined.

In the current study we did not find any significant relationships between measures of cognitive performance and measures of neural variability despite the use of several different tasks with distinct stimuli, structures, and attentional/cognitive demands. Note that this disappointing result was apparent across two independent experimental sessions. This suggests that the relationship between neural variability and cognitive performance is not as strong and clear as previously suggested (Garrett et al., 2013), in contrast to the relationship with perceptual performance, which has been reproduced by several labs using different neuroimaging and analysis techniques (Schurger et al., 2010, 2015; Xue et al., 2010; Arazi et al., 2017). Regardless of the precise relationship between neural variability magnitudes and behavioral performance, the main contribution of this study is in demonstrating that neural variability magnitudes differ across individuals in a reliable manner. These individual differences in neural variability are likely to constrain individual behavioral performance at some level, whether only with respect to perceptual performance or also in other behavioral domains.

### Signal strength and trial-by-trial variability

Previous electrophysiology studies have demonstrated that trial-by-trial variability scales with the mean amplitude of the examined neural responses (Shadlen and Newsome, 1994). To examine trial-by-trial variability independently of the mean response most electrophysiology studies, therefore, use the Fano Factor or the coefficient of variation (CV), which normalize trial-by-trial variability by the mean response (Churchland et al., 2010; Goris et al., 2014). Animal studies, however, have rarely examined the behavioral impact of neural variability magnitudes.

In contrast, human neuroimaging studies that have examined the relationship between response variability and behavior using fMRI and EEG have rarely reported CV (Garrett et al., 2013, 2015; He and Zempel, 2013; Gonen-Yaacovi et al., 2016; Arazi et al., 2017). Nevertheless, to relate our findings to both literatures, we carried out our analyses once using trial-by-trial variability measures and again using the CV measure. We found almost equivalent results in both cases, which revealed that large between-subject differences in variability magnitudes are consistent across experimental sessions and tasks even when normalizing trial-by-trial variability by the mean EEG response. Consistent differences in neural variability magnitudes across subjects are, therefore, likely to reflect differences in underlying physiological mechanisms that are specific to the variability of neural activity rather than the strength of neural activity.

### Measurement noise

Measures of trial-by-trial neural variability may be biased by subject-specific measurement noise of non-neural origin. We examined two potential sources of non-neural variability in our study: eye-gaze variability (indicative of the stability of fixation across trials) and trial-by-trial variability in electrode offset (indicative of the stability of the EEG recording). We did not find any significant correlation between electrode-offset variability or gaze-position variability and neuronal measures of variability. Furthermore, regressing out individual magnitudes of electrode offset variability or gaze position variability did not alter the results. These additional analyses demonstrate that the individual magnitudes of trial-by-trial variability were not associated with these potential sources non-neural measurement noise. With that said, additional studies examining the consistency of individual neural variability magnitudes across different neuroimaging techniques (e.g., fMRI and EEG) would be necessary for demonstrating the potential robustness of these findings across techniques with different types/sources of measurement noise.

### Conclusions and future directions

While neural variability is to some degree under flexible control of attention and neuromodulation, our results demonstrate that individual differences in neural variability magnitudes are remarkably consistent across distinct tasks and over long periods of time. We, therefore, propose that neural variability magnitudes represent mostly stable between-subject differences in fundamental neural characteristics that were forged by genetics and environmental exposures during early development. These differences are likely to predispose individual subjects to exhibit distinct behavioral capabilities. Revealing how neural variability magnitudes develop during childhood and how they may be manipulated in adulthood are likely to be of great interest for further basic and clinical research (Dinstein et al., 2015).

## Acknowledgments

This study was supported by ISF grant 961/14 (ID) and ministry of immigrant absorption fellowship (GGY). The authors declare no competing financial interests.

## Author contributions

A.A., G.G.Y and I.D. design the experiment. A.A. and G.G.Y performed the experiment. A.A analysed the data. A.A, and I.D wrote the paper.

